# Internalisation and biological activity of nucleic acids delivering cell-penetrating peptide nanoparticles is controlled by the biomolecular corona

**DOI:** 10.1101/2021.03.26.437157

**Authors:** Annely Lorents, Maria Maloverjan, Kärt Padari, Margus Pooga

**Affiliations:** Institute of Molecular and Cell Biology, University of Tartu, Tartu, Estonia; Institute of Technology, University of Tartu, Tartu, Estonia

**Keywords:** Cell-penetrating peptide, nucleic acid delivery, CPP nanoparticles, protein corona

## Abstract

Nucleic acid molecules can be transferred into cells to alter gene expression and, thus, alleviate certain pathological conditions. Cell-penetrating peptides (CPPs) are vectors that can be used for transfecting nucleic acids as well as many other compounds. CPPs associate nucleic acids non-covalently, forming stable nanoparticles and providing efficient transfection of cells *in vitro*. However, *in vivo*, expected efficiency is achieved only in rare cases. One of the reasons for this discrepancy is formation of protein corona around nanoparticles, once they are exposed to a biological environment, e.g. blood stream. In this study, we compared CPP-nucleic acid nanoparticles formed in the presence of bovine, murine and human serum. We used Western blot and mass-spectrometry to identify the major constituents of protein corona forming around nanoparticles, showing that proteins involved in transport, haemostasis and complement system are its major components. We investigated physical features of nanoparticles, and measured their biological efficiency in splice-correction assay. We showed that protein corona constituents might alter the fate of nanoparticles *in vivo*, e.g. by subjecting them to phagocytosis. We demonstrated that composition of protein corona of nanoparticles is species-specific that leads to dissimilar transfection efficiency and should be taken into account while developing delivery systems for nucleic acids.

## Introduction

Development of functionalised nano-tools for biomedical research and therapy is explosively progressing field and various strategies are applied for refining NPs, from both the vector and the biologically active cargo side.^1–4^ Among highly promising approaches is application of CPPs^5–7^ as efficient carrier vectors for the delivery of different functional nucleic acid (NA) molecules, e.g. plasmid DNA, mRNA, siRNA, miRNA, splicing correcting oligonucleotides etc.^8^ Once transfected, these NAs can modulate the flow of genetic information in the cells and, consequently, alter cellular responses in target sites.^9^ CPPs are able to condense NA by electrostatic complexing after simple mixing step and effectively transport compacted NA into the cells, mostly by harnessing various endocytic pathways. Delivery of NAs with CPPs *in vitro* results in efficient and homogeneous uptake of the complexes by cells and in a functional effect of NA, e.g. expression of exogenous gene from pDNA or silencing of targeted gene by siRNA. However, application of the same delivery system in animal models *in vivo* has usually yielded much lower efficiencies so far,^4,10^ which is also characteristic for lipid-based^11^ and other types of nanoparticles.^12,13^ Still, encouragingly, several break-through studies regarding applicability of CPP-mediated transport *in vivo* have been published recently.^14–16^

Nowadays, it is well established that various proteins rapidly coat the NPs engineered for drug delivery after the contact with biological fluids (e.g. after contact with bloodstream in case of intravenous administration).^17–19^ Formation of a protein corona provides the pristine NPs with a new biomolecular interface for interaction with blood cells and lining endothelium, and for detection by specialised phagocyte cells with subsequent elimination.^20,21^ The components of protein corona rather than the nanoparticle *per se* are typically recognized by receptors on cell surface. For example, in the case of DNA-lipid nanoparticles, vitronectin enriched on particles associates with αvβ3 integrins on cancer cells.^22^ Analogously, for targeting to hepatocytes, interaction with apolipoproteins is required for some siRNA lipoplexes.^23,24^ Remarkably, the liver tropism of the first siRNA drug, Onpattro, is based on the ability of lipid nanoparticle to recruit into its PC apolipoprotein E that enables specific hepatocyte targeting.^11,25,26^ Recently, the preformed protein corona was harnessed to convert nanoparticles targetable to tumour cells. For that, particles were coated with tumour-specific antibodies and human serum albumin that helps to prevent phagocytosis of the particles.^27–29^

On the other hand, protein corona can interfere with targeted delivery by masking the ligand on the nanoparticle surface that specifically interacts with endocytosis triggering receptors.^30,31^ The composition and properties of forming protein corona highly depend on one hand on the type and properties of nanoparticles, like composition, size, charge, hydrophobicity etc.,^10,32–35^ and on the other, on environmental factors, like temperature,^36,37^ incubation time^17,38^ and, especially, on the biofluid type,^39,40,41^ origin and disease^42,43^. In general, protein corona on the smaller nanoparticles is thinner and less dense than on the larger ones.^44,45^ In addition, cationic nanoparticles are cleared from circulation much faster than neutral or anionic counterparts.^46^ Thus, one of the potential obstacles for cellular uptake of CPP-nucleic acid nanoparticles *in vivo* might be their inefficient association with cells due to the formation of protein corona on the surface of particles after introduction into physiological environment.^47^

Condensation of differently sized nucleic acid molecules to nanoparticles using various cell-penetrating peptides has been well documented.^48–51^ Although the properties of such nanoparticles might significantly change after formation of protein corona on its surface, the composition of protein corona and its impact have not been studied so far. Only the increase of the nanoparticle’s size in the presence of serum^48,50,52–54^ and the decrease in protein quantity associating with particle upon reducing CPP/nucleic acid molar ratio have been reported.^55^

In this study, we characterised the composition and effect of the protein corona that formed on the surface of CPP-nucleic acid nanoparticles. Analysed particles contained plasmid DNA (pDNA) or splice-correcting oligonucleotide (SCO) that were non-covalently complexed with three CPPs: PepFect 14 (PF14), PF14 modified with a 22-carbon fatty acid tail (C22-PF14)^56^ or NickFect 55 (NF55).^57^ We studied the formation of protein corona on different nanoparticles in bovine serum as a representative of cell culture systems, in murine serum as a typical animal model used in the development and testing of such nanoparticles, and in human serum as a representative of the actual target for drug development (Schema 1). We observed that dissimilar protein corona forming in different sera translates to unequal uptake of studied nanoparticles by cultured cells, and varying biological response to nucleic acid cargo. Our results emphasize that, as expected, the features of protein corona on CPP-nucleic acid nanoparticles vary in different animal models and cell culturing conditions, and these might represent the underlying cause for discrepant results obtained in *in vitro* and *in vivo* systems respectively.

**Schema 1.**
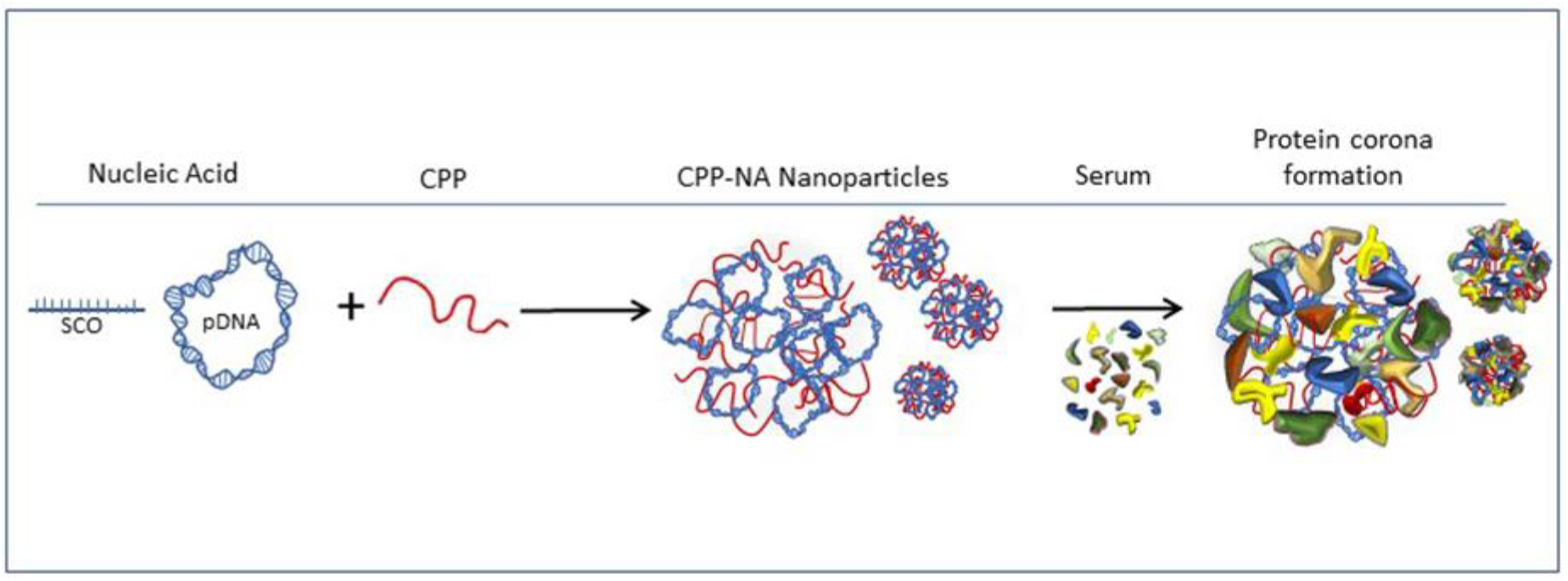
Condensation of nucleic acid (NA) to nanoparticles by cell-penetrating peptide (CPP), and formation of protein corona on nanoparticles in serum containing media.

## Results and Discussion

### Formation of protein corona on CPP-pDNA nanoparticles depends on used serum

PepFect and NickFect series of CPP were designed for delivering into cells nucleic acid molecules with different biological activity. As expected, all three used CPPs condensed nucleic acids to nanoparticles with 100-200 nm diameter (see Table 1 for used components, Table 2 for characteristics of formed nanoparticles, and SFig. 1 for DLS analysis) at optimal CPP/nucleic acid ratios.^49^ The highest transfection efficiency with these CPPs has been achieved at higher charge ratio (N/P ratio 2) with pDNA, whereas SCO requires lower excess of peptide (N/P ratio 1.5) for transduction into cells, which also leads to different zeta potential of formed nanoparticles.

**Table 1.**
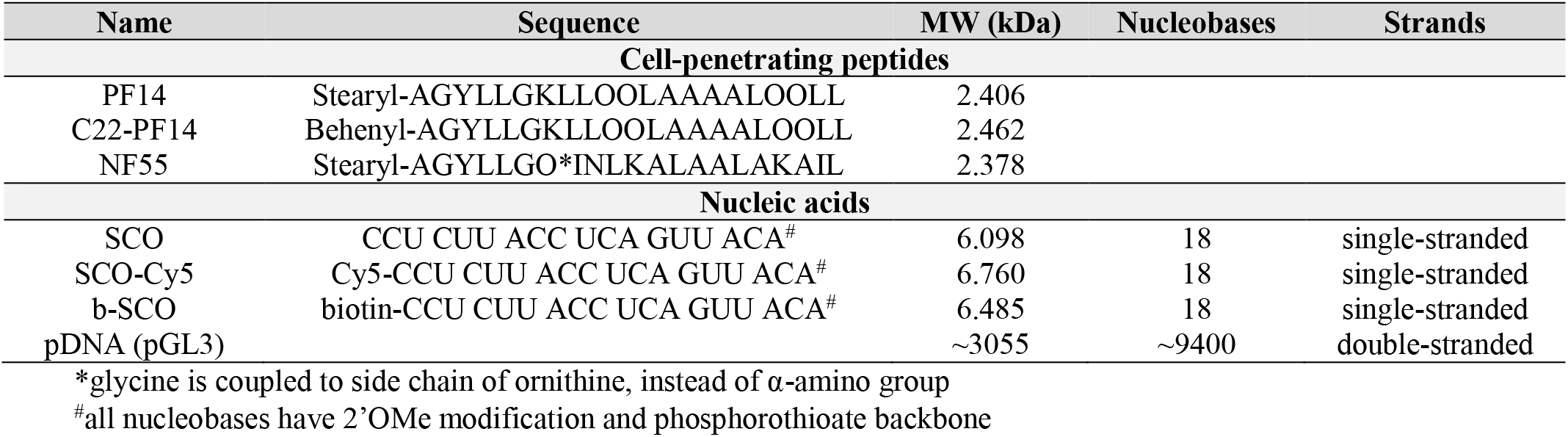
Characteristics of cell-penetrating peptides (CPP) and nucleic acids used in nanoparticles.

**Table 2.**
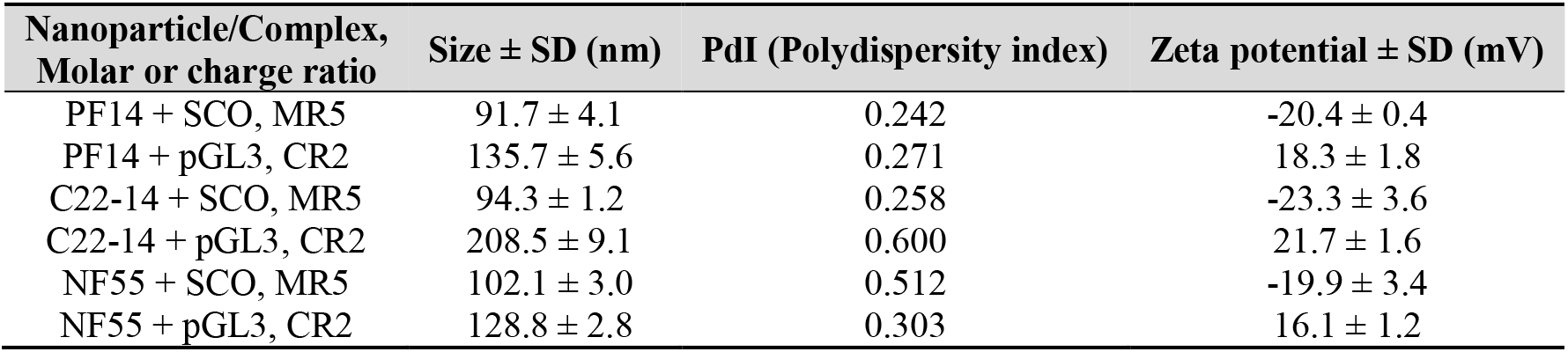
Hydrodynamic diameter of CPP-nucleic acid nanoparticles in water, and zeta potential in 1 mM NaCl.

To assess the formation of protein corona around the CPP-nucleic acid nanoparticles, we captured nanoparticles consisting of biotinylated pDNA and studied CPPs to streptavidin-paramagnetic beads (Schema 2). After incubation with 50% serum, known to resemble the *in vivo* conditions,^58^ and washing, we collected the beads with bound nanoparticles-, subjected the samples to gel electrophoresis to separate adsorbed proteins and visualised proteins by silver staining. Our results demonstrated that a highly complex protein corona formed around nanoparticles already after 5 min incubation with both, bovine and murine serum (Fig. 1).^19^ After 60 min incubation with serum, protein profiles were comparable to the ones observed after 5 min, however, some bands were more intense after longer incubation time, consistent with gradual adsorption of proteins.^17,38,59^ Most prominent band in all cases was a protein with approximate size of 70 kDa and probably represents serum albumin as it constitutes about half of serum proteins, and is abundant in protein corona of all types of nanoparticles.^11,60^ Even though, the differences in protein profiles adsorbed to nanoparticles formed with PF14 or NF55 were negligible according to gel electrophoresis results. It was clearly observable that nanoparticles incubated in bovine (Fig. 1a) or murine serum (Fig. 1b) captured a different protein ensemble. This suggests that not only the physical-chemical properties of nanoparticles play a role in formation of protein corona, but the nature of the corona can also depend on the studied animal. We assume this is the underlying cause why CPP-nucleic acid nanoparticles that are efficient in one model system, e.g. *in vitro* in cell culture, may fail in another, e.g. *in vivo*. This discrepancy should also be considered while developing and testing such nanoparticles for efficient delivery in differing milieu. Analogous, species-selective association with plasma proteins was recently reported for antisense oligonucleotides.^60^

**Figure 1.**
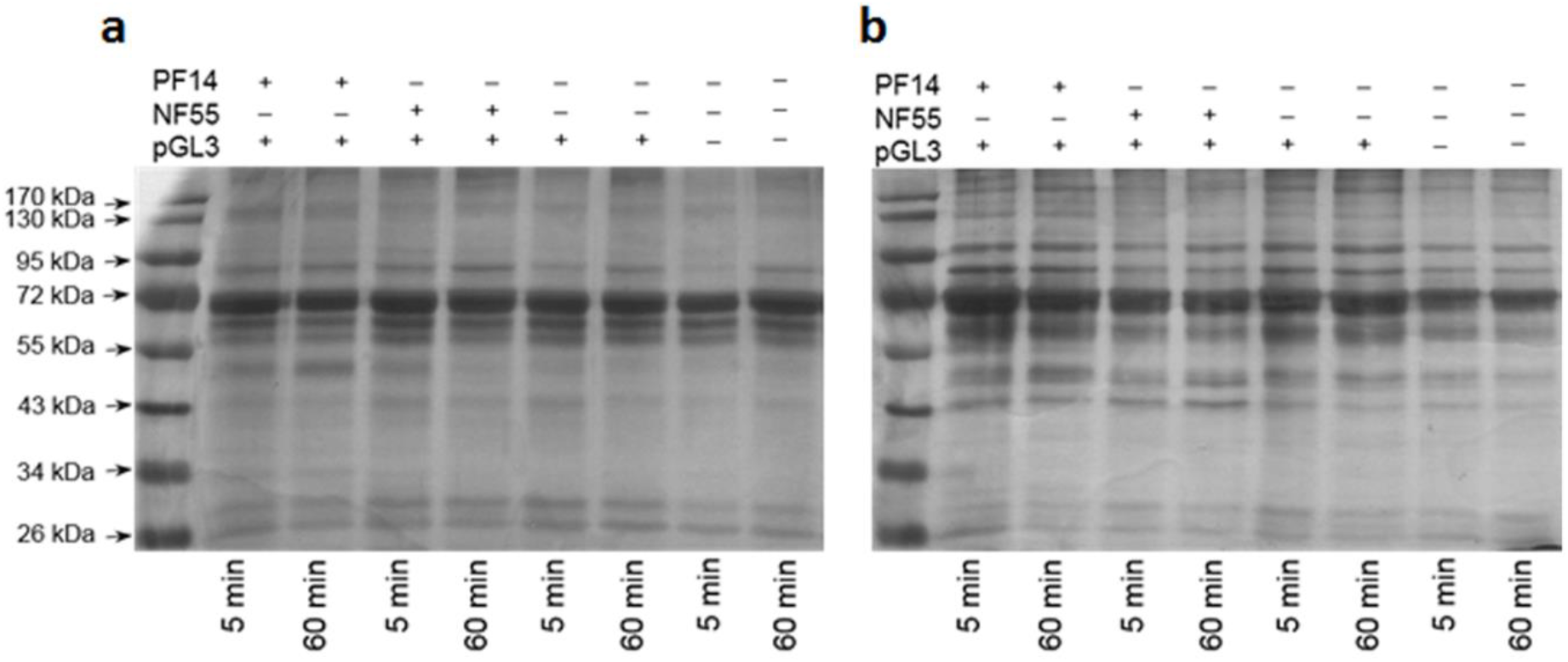
Adsorption of proteins on the CPP-plasmid DNA nanoparticles. CPP-pGL3 nanoparticles or pGL3 alone were immobilised on streptavidin-superparamagnetic beads, and incubated in foetal bovine serum (**a**) or murine serum (**b**) for 5 or 60 minutes at room temperature. After pull-down of formed complexes, proteins were separated on 9% acrylamide gel and visualised by silver staining. One representative of three independent experiments is presented.

**Schema 2.**
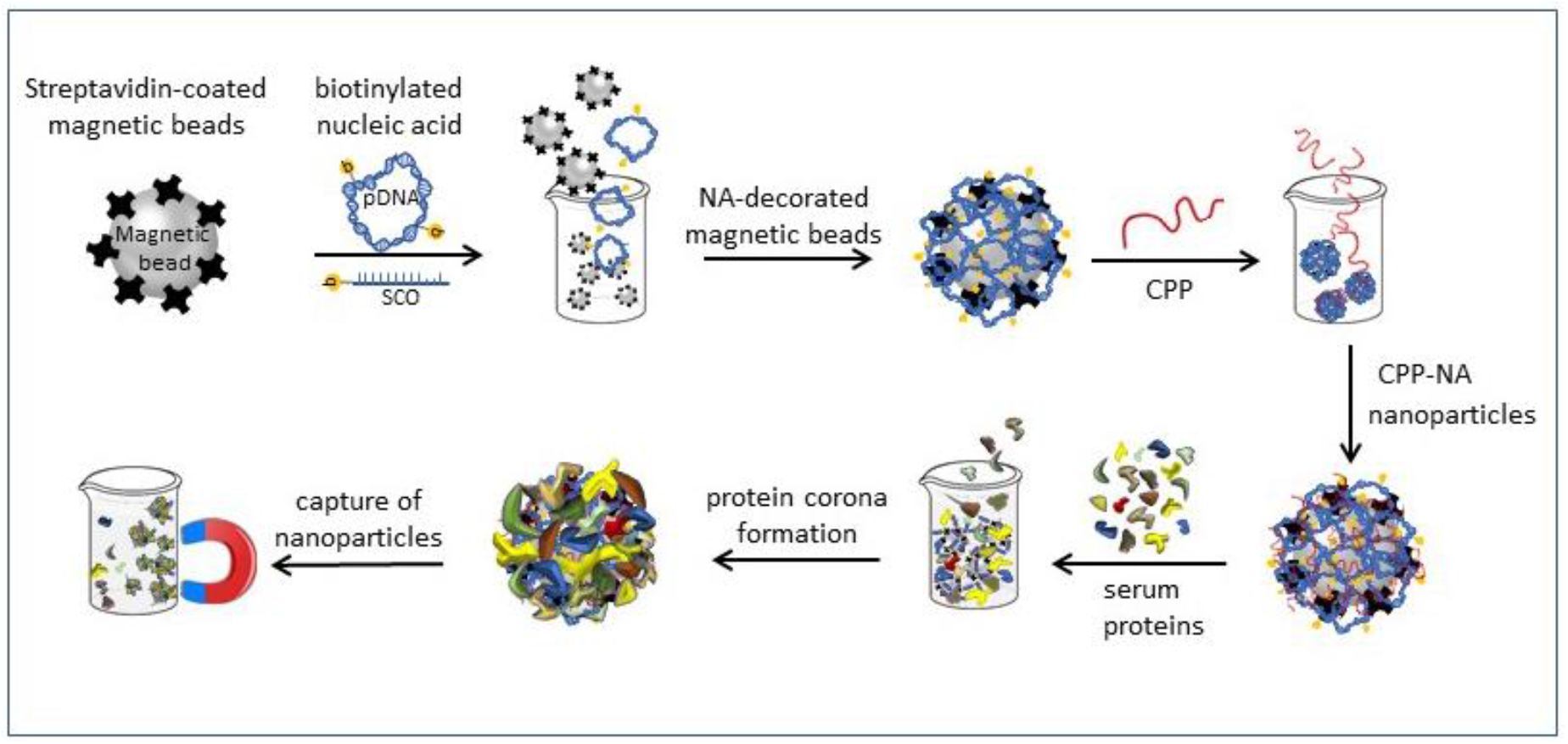
Condensation of biotinylated nucleic acid (NA, pDNA or SCO) with cell-penetrating peptide (CPP) into nanoparticles, and formation of protein corona on the nanoparticles on the surface of magnetic beads. Capture of nanoparticles and protein corona immobilised on magnetic beads from protein containing solution by magnet for further analysis of proteins in corona.

### Nanoparticles may aggregate in serum containing environment

Typically, the characteristics of CPP-nucleic acid nanoparticles like size, zeta potential and morphology are analysed in media used for assembly of particles, e.g. in diluted buffer solution or water (Table 2).^49,50,52,61^ Therefore, next we attempted visualisation of protein corona on CPP-nucleic acid nanoparticles to assess, whether the morphology or aggregation of particles could change upon contact with biologically relevant environment that has been demonstrated earlier for other types of nanoparticles.^51,62^ We associated PF14 with biotin-labelled SCO and tagged the formed complexes with 10 nm gold particles via neutravidin (black dots in Fig. 2)^63^ for facilitating detection by transmission electron microscopy (TEM). In water, mostly single nanoparticles of characteristic shape^49^ formed and some had gathered to small clusters as verified by TEM in specimens with negative staining with uranyl acetate (Fig. 2a). Analogously, in unstained specimen of PF14-SCO nanoparticles, the gold label was randomly distributed, and due to low electron density, the localisation of nanoparticles can only be assigned based on localisation of the gold label (Fig. 2b). However, after incubation with serum, single or slightly clustered PF14-SCO nanoparticles had formed conglomerates with size of several hundred nanometres. Furthermore, the adsorbed proteins had increased the electron density of clusters to level detectable by TEM even in unstained specimens (Fig. 2d-f), corroborating formation of extensive protein corona. Still, the electron density of adsorbed proteins was not sufficient for analysing the actual or detailed morphology of protein corona formed around nanoparticles. The shape of conglomerates induced by the formation of protein corona resembled rather a cloud of loosely associated proteins than a regular layer with uniform thickness as was also observed earlier for other nanoparticles.^33^

**Figure 2.**
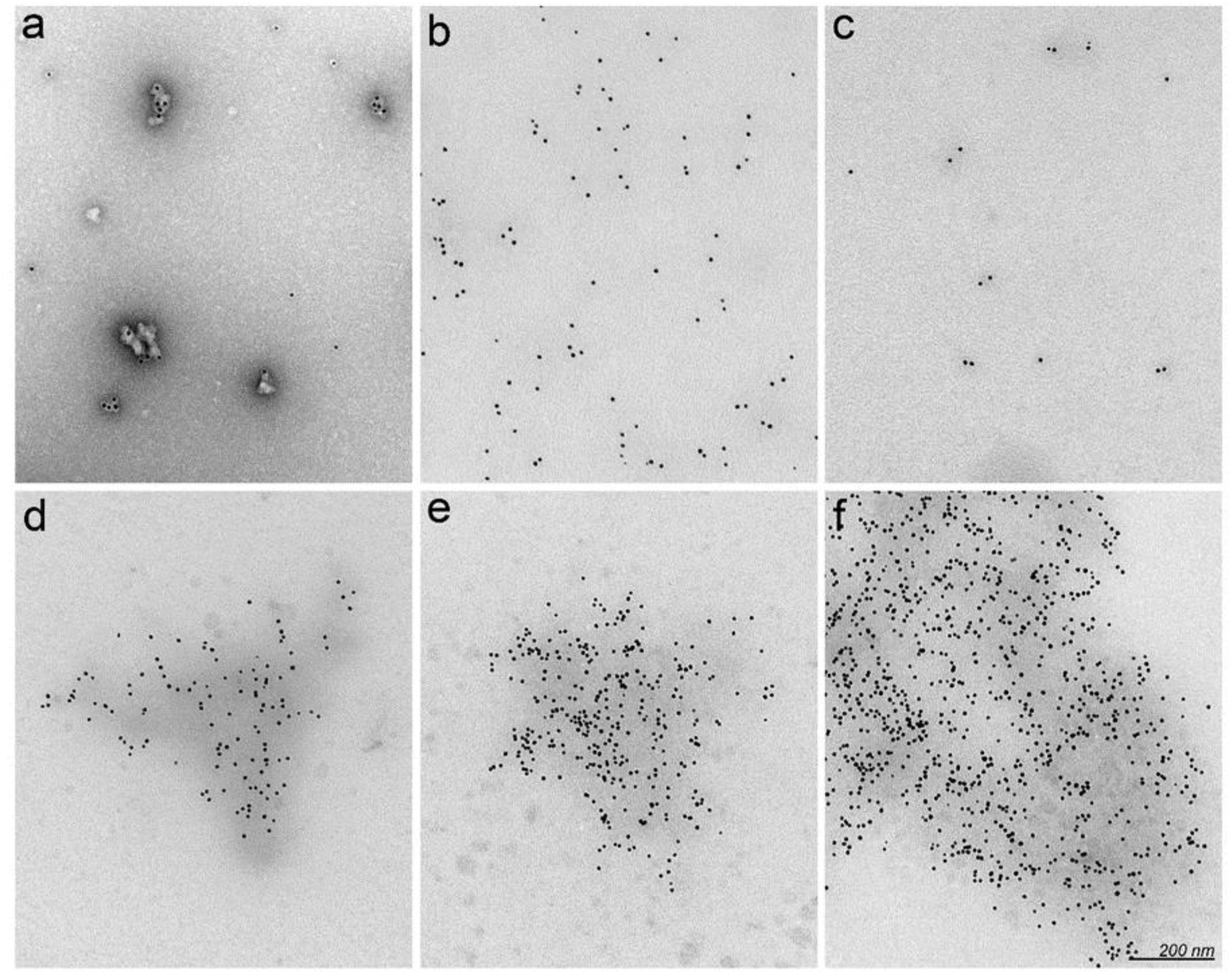
Visualisation of PF14-SCO nanoparticles and protein corona by transmission electron microscopy (TEM). Complexes of PF14 and biotin-SCO labelled with colloidal gold of 10 nm diameter (black dots) were formed in water at PF14/SCO molar ratio 5. PF14-SCO complexes without protein corona negatively stained with uranyl acetate on TEM grid (**a**), and in unstained specimen (**b**). Biotin-SCO labelled with colloidal gold and stained with uranyl acetate (**c**). Protein corona forming in 50% bovine (**d**), murine (**e**) or human serum (**f**) clusters PF14-SCO nanoparticles to conglomerates with electron dense background. Scale bar: 200 nm. Representative images from three independent experiments are presented.

Unfortunately, negative staining that might resolve the morphology of protein corona on nanoparticles, produced abundant artefactual structures with serum proteins that interfered with the visualisation of protein corona as has been reported earlier.^64^ Cationic delivery systems for nucleic acids often reveal pulmonary toxicity^65^ that has been associated with clogging of small lung capillaries.^66^ Aggregation of nanoparticles by serum proteins that we observe in TEM experiment might exacerbate this even more. However, aggregation of PF14-SCO nanoparticles in real *in vivo* conditions after administration is probably less extensive than in static conditions applied here due to dynamic flow and shear stress that strongly influence the structure and composition of biomolecular corona.^67,68^

### Proteins involved in transport and haemostasis are major constituents of protein corona

In order to dissect the composition of protein corona that forms in animal sera in detail, we captured biotin-tagged nanoparticles on streptavidin-coated magnetic beads, incubated with serum, and performed mass-spectrometry analysis of absorbed proteins. First, we evaluated formation of protein corona around “naked” streptavidin beads and confirmed that it was negligible compared to pDNA-carrying streptavidin-beads or beads that carried CPP-pDNA nanoparticles (Table 3). Next, we identified proteins bound to pGL3 alone or pGL3 complexed with CPPs. As expected, serum albumin that is known also to associate tightly with “naked” cationic CPPs^69^ showed the highest abundance in all samples after incubation of nanoparticles with bovine (Table 3), murine (Table 4), or human serum (Table 5).

**Table 3.** Mass-spectrometry analysis of proteins bound to CPP-pDNA nanoparticles in bovine serum at higher level. PF14-pDNA nanoparticles or pDNA alone were coupled to streptavidin-coated superparamagnetic beads and incubated with bovine serum for 30 minutes at room temperature. After pull-down of beads with magnet and washes, the proteins of PC were digested with trypsin and analysed by mass-spectrometry. Overall number of identified peptides of a protein group is presented. Fifteen most abundant proteins in protein corona are listed. Data represents two independent experiments and three technical replicates of each with ±SEM, compared to b-pGL3. \**p-value* < 0.05, \*\**p-value* < 0.005, \*\*\**p-value* < 0.0005.

| Protein group | Beads | + b-pGL3 | + PF14 + b-pGL3 |
| --- | --- | --- | --- |
| Serum albumin | 25.5 ± 0.5 | 67.5 ± 1.5 | 71.5 ± 1.5 |
| α-2-HS-glycoprotein | 7.0 ± 0.0 | 17.0 ± 0.0 | 19.5 ± 0.5 |
| Apolipoprotein A-II | 3.5 ± 0.5 | 9.5 ± 0.5 | 10.0 ± 0.0 |
| α-1-antiproteinase | 7.0 ± 0.0 | 27.0 ± 1.0 | 27.5 ± 0.5 |
| Apolipoprotein A-I | 6.5 ± 0.5 | 28.0 ± 0.0 | 35.0 ± 0.0*** |
| Serotransferrin | 5.5 ± 0.5 | 52.5 ± 7.5 | 61.5 ± 2.5 |
| Haemoglobin subunit $\alpha$ | 4.0 ± 0.0 | 9.0 ± 0.0 | 10.0 ± 0.0** |
| Haemoglobin foetal subunit $\beta$ | 4.0 ± 0.0 | 15.0 ± 0.0 | 16.5 ± 0.5 |
| α-2-macroglobulin | 4.5 ± 0.5 | 59.0 ± 7.0 | 69.5 ± 0.5 |
| Fetuin-B | 0.0 ± 0.0 | 12.0 ± 0.0 | 13.0 ± 0.0** |
| Inter-alpha-trypsin inhibitor heavy chain H2 | 6.0 ± 0.0 | 25.5 ± 0.5 | 34.5 ± 0.5** |
| Vitamin D-binding protein | 3.0 ± 1.0 | 25.5 ± 3.5 | 27.5 ± 0.5 |
| Prothrombin | 5.0 ± 1.0 | 13.0 ± 1.0 | 34.0 ± 1.0* |
| Complement component 3 | 6.5 ± 0.5 | 35.0 ± 4.0 | 62.5 ± 0.5 |
| α-1-acid glycoprotein | 0.0 ± 0.0 | 7.0 ± 1.0 | 10.0 ± 0.0 |

**Table 4.**
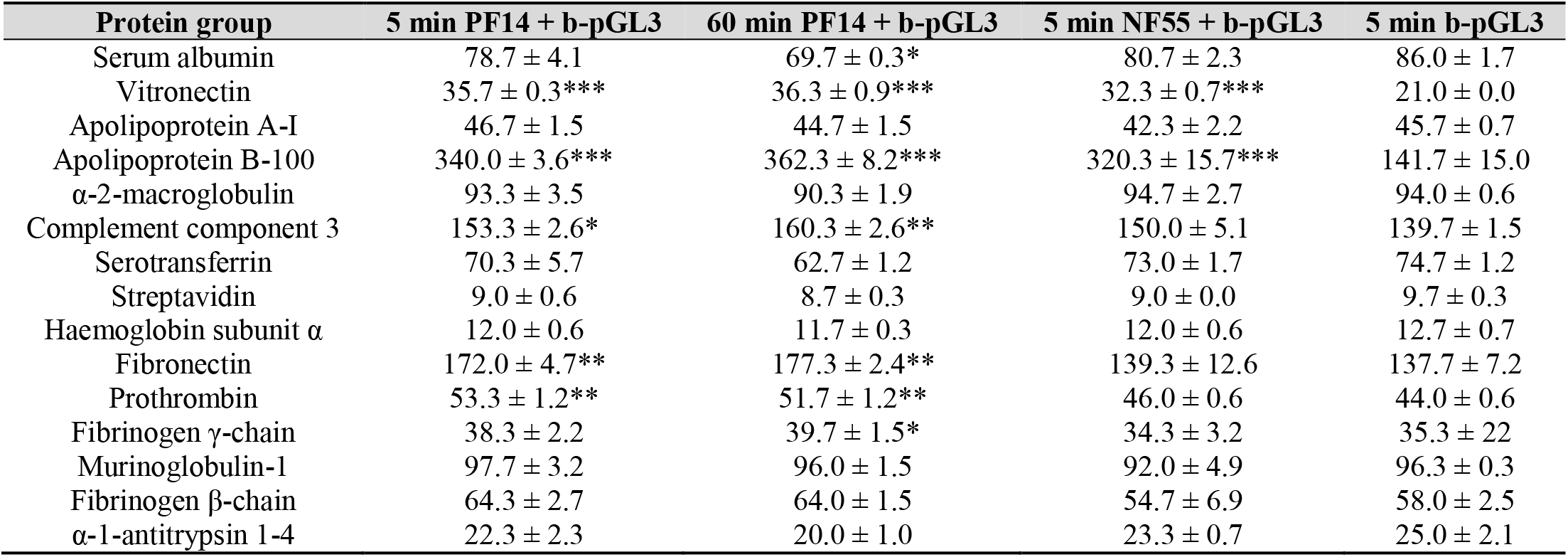
Mass-spectrometry analysis of proteins bound to CPP-pDNA nanoparticles from murine serum at higher level. Fifteen most abundant proteins in protein corona are listed. Data represents two independent experiments and three technical replicates of each with ±SEM, compared to b-pGL3. \**p-value* < 0.05, \*\**p-value* < 0.005, \*\*\**p-value* < 0.0005.

**Table 5.** Mass-spectrometry analysis of proteins bound to CPP-pDNA nanoparticles from human serum at higher level. Fifteen most abundant proteins in protein corona are listed. Data represents two independent experiments and three technical replicates of each with ±SEM, compared to b-pGL3. \**p-value* < 0.05, \*\**p-value* < 0.005, \*\*\**p-value* < 0.0005.

| Protein group | b-pGL3 | PF14 + b-pGL3 |
| --- | --- | --- |
| Serum albumin | 44.5 ± 0.5 | 91.5 ± 0.5*** |
| Serotransferrin | 15.5 ± 0.5 | 76.0 ± 0.0** |
| α-2-macroglobulin | 20.0 ± 1.0 | 89.5 ± 1.5* |
| α-1-antitrypsin | 23.0 ± 0.0 | 33.5 ± 1.5 |
| Apolipoprotein A-I | 15.5 ± 0.5 | 30.0 ± 0.0* |
| Haptoglobin | 16.0 ± 1.0 | 27.5 ± 0.5 |
| Complement component 3 | 16.0 ± 1.0 | 119.5 ± 0.5** |
| Ig γ-1 chain C region | 9.0 ± 1.0 | 15.0 ± 0.0 |
| Ig γ-3 chain C region | 6. ± 0.5 | 17.5 ± 0.5 |
| Ig κ-chain C region | 7.0 ± 0.0 | 12.0 ± 0.0 |
| α-1-acid glycoprotein 1 | 5.0 ± 0.0 | 12.0 ± 0.0*** |
| Ig α-1 chain C region | 7.0 ± 1.0 | 20.0 ± 0.0* |
| Complement component 4-B | 4.0 ± 0.0 | 78.0 ± 1.0** |
| Hemopexin | 0.0 ± 0.0 | 25.0 ± 0.0*** |
| Apolipoprotein A-II | 5.5 ± 0.5 | 8.0 ± 0.0 |

Adsorption of albumin, the most abundant protein in sera, and prominent constituent in protein corona of all nanoparticles,^70^ grants certain advantages to nanoparticles. Coverage with albumin has been shown to reduce the lysosomal degradation of nanoparticles in endothelial cells and, therefore, extend the circulation time of particles.^47,71^ Furthermore, albumin facilitates transport of nanoparticles from the site of injection to peripheral tissues,^60^ endocytosis^72^ and transcytosis across capillary endothelium.^73^ In addition, other plasma proteins involved in binding, transport, and mediating availability of various biomolecules in the bloodstream [e.g. apolipoproteins, α-2-HS-glycoprotein (also known as fetuin-A), serotransferrin etc.]; and proteins involved in blood coagulation and haemostasis (e.g. α-2-macroglobulin, prothrombin, fibrinogen, α-1-antitrypsin etc.) were identified in protein corona of CPP-nucleic acid nanoparticles. Moreover, contact activation pathway of coagulation can be activated by artificial negatively charged substances, including oligonucleotides, “naked” pDNA and anionic nanoparticles that may explain the presence of coagulation proteins in protein corona of CPP-pDNA nanoparticles.^74,75^

Fetuin found in the protein corona has been demonstrated earlier to trigger internalisation of nanoparticles, e.g. the uptake of nanospheres by Kupffer cells through the scavenger receptors.^76^ Furthermore, addition of fetuin into culture medium increased cellular uptake of PF14-SCO nanoparticles and splicing correction by SCO in HeLa pLuc705 cells in SR-dependent manner.^52^ Interestingly, in case of human serum, PF14-pDNA nanoparticles demonstrated higher adsorption of most proteins compared to pDNA alone (Table 5), further emphasizing marked differences in composition of protein corona between sera of various mammals. In addition to fetuin, which marks nanoparticles for recognition by macrophages via scavenger receptors, also α-2-macroglobulin that is abundant in protein corona of CPP-nucleic acid nanoparticles (Tables 3–5) might dampen the effect of injected pharmaceuticals, like oligonucleotides, targeting these to elimination as found earlier.^77^

To sum up, serum albumin was the most abundant protein in protein corona forming around CPP-nucleic acid nanoparticles in all studied sera: from calf, mouse and human. Still, the protein composition varied extensively between different sera and each protein corona showed individual pattern.

In addition, the functioning of proteins forming corona around PF14-pDNA nanoparticles in murine serum is in good concordance with the recent reports about the behaviour and targeting^78^ of these nanoparticles *in vivo*. Accordingly, high level of alpha 1-antitrypsin in protein corona of PF14-pDNA nanoparticles (Table 3–5) correlates well with accumulation of nanoparticles in lungs of mice.^79^ It was also shown, that accumulation of apolipoprotein B-100,^80^ and complement component 3^78^ in protein corona targets nanoparticles to liver. In both these organs high expression of luciferase from pGL3 plasmid was observed recently, whereas only low amounts of these nanoparticles were targeted to spleen, kidney, and heart.^55,81^

### Adsorption of complement component 3 depends on nanoparticle characteristics

Besides carrier proteins found in blood and proteins involved in haemostasis, mass spectrometry analysis also revealed the high level of complement factors in the protein corona of studied nanoparticles, especially of complement component 3 (C3) that associated with complexes most abundantly in murine serum. Nanoparticles are cleared from the bloodstream by opsonisation that is induced by antibodies and complement proteins associating with exogenous particles; and C3 plays a central role in the activation of complement system via all three pathways,^82–84^ and, thus, opsonisation.^83,85^ Therefore, we further focussed on the level of C3 in the corona formed under different conditions. I.e. in the plasma that contains active complement factors as compared with serum, where these were inactivated by heating. As previously, pDNA and SCO were complexed with CPPs, incubated with serum or plasma. After pull-down with streptavidin-coated magnetic beads, C3 levels in protein corona were analysed by Western blot using an antibody to the C-terminus of C3 that recognizes C3 α-chain and also fragment 2 of α-chain that corresponds to C3d.^86^ First, we evaluated adsorption of C3 to pDNA complexed with PF14, NF55, and C22-PF14. After 60 min incubation with murine serum (Fig. 3a, b; SFig. 2) or murine plasma (Fig. 3c, d; SFig. 1, 2) no significant difference in C3 levels was detected between nanoparticles formed with different CPPs. However, pDNA alone and NF55-pDNA complexes showed tendency to adsorb higher amount of complement proteins as compared to nanoparticles containing PF14 and C22-PF14 (Fig. 3), although this difference was not statistically significant. In addition, our results showed that after 60 min incubation, the level of C3 α-chain fragment 2 bound to PF14-pDNA nanoparticles was lower than after incubation for 60 min that was in line with time-dependent adsorption of proteins on nanoparticles.^17,59^ C3 α-chain and its fragment 2 associated at significantly higher level to beads covered with “naked” pDNA in murine serum (Fig. 3a) and plasma (Fig. 3c) when compared to PF14-containing nanoparticles, suggesting that condensation of pDNA to nanoparticles with PF14 prolongs its circulation *in vivo*. Still, higher accumulation of C3 and its fragments into protein corona in murine plasma (Fig. 3d) compared to serum (Fig. 3b) favours elimination of CPP-pDNA nanoparticles from circulation in mouse *in vivo*, where the complement system is active, compared to *in vitro* conditions, where cells remain exposed to higher amounts of complexes in serum-containing medium.

**Figure 3.**
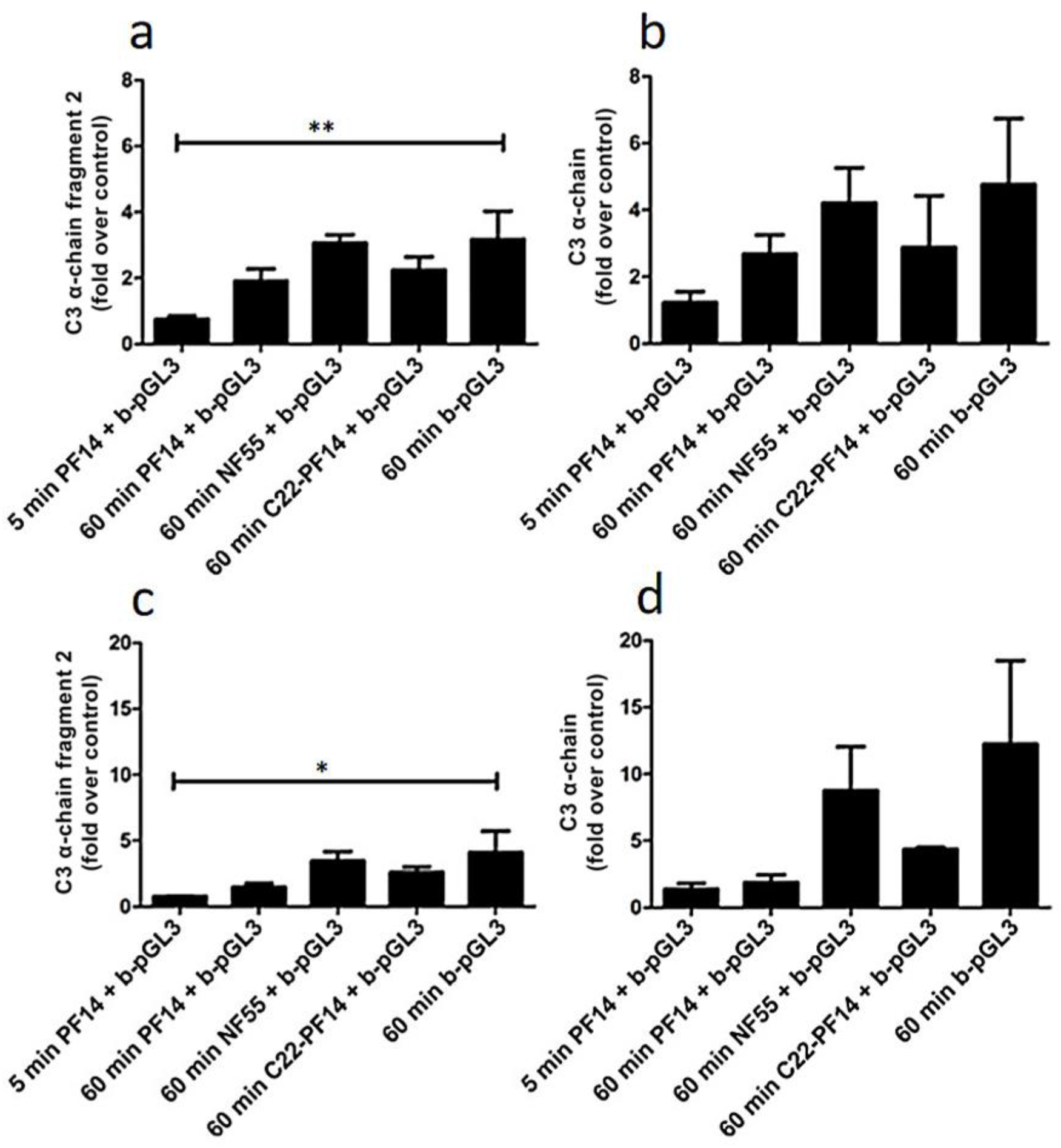
Level of complement component 3 (C3) in protein corona of plasmid DNA and CPP-pDNA nanoparticles. CPP-pGL3 nanoparticles or pGL3 alone were immobilized on streptavidin-superparamagnetic beads, or beads were left untreated (control) before incubation with murine serum (**a, b**) or murine plasma (**c, d**) for 5 or 60 minutes at room temperature. After pull-down, bound C3 subunits and its fragments were quantified by Western blot. Data represents at least three independent experiments, ±SEM, analysed by one-way ANOVA with Dunnett’s test, \**p-value* < 0.05, \*\**p-value* < 0.005.

Next, we assessed inclusion of C3 in protein corona of SCO-containing nanoparticles. Upon incubation of PF14-SCO nanoparticles with murine serum, no significant difference was observed between 5 min and 60 min incubation, and the level of C3 proteins in protein corona stayed at a similar level with SCO alone (Fig. 4a, b, SFig. 4). However, incubation of NF55-SCO nanoparticles in murine serum for 60 min resulted in a significantly higher amount of C3 proteins in the corona compared to SCO alone. That may be caused by higher hydrophobicity of NF55^81^ or its complexes with SCO compared to PF14, and higher absorption of proteins, including C3,^87^ on hydrophobic NPs.^88^ In case of murine plasma, however, no significant differences were observed between any nanoparticles tested and the amount of C3 proteins adsorbed to nanoparticles was rather equal (Fig. 4c, d; SFig. 5).

**Figure 4.**
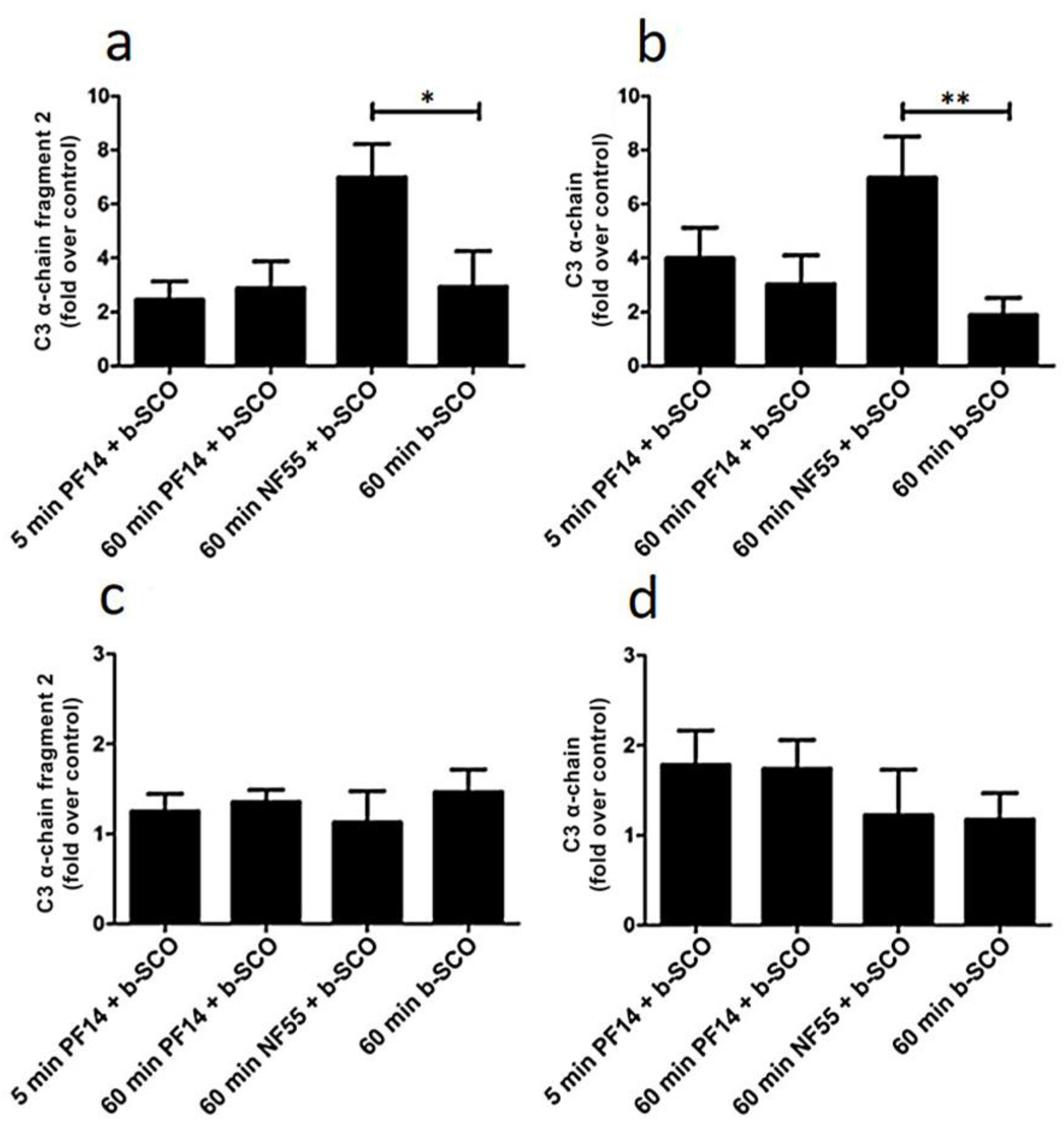
Level of complement component 3 (C3) in protein corona of CPP-SCO nanoparticles. CPP-SCO nanoparticles or SCO alone were bound to streptavidin on superparamagnetic beads or beads were left untreated (control) before incubation with murine serum (**a, b**) or murine plasma (**c, d**) for 5 or 60 minutes at room temperature. After pull-down, bound C3 and its fragments were quantified by Western blot. Data represents at least three independent experiments, ±SEM, analysed with one-way ANOVA with Dunnett’s test, \**p-value* < 0.05, \*\**p-value* < 0.005.

The second generation CPPs of PepFect and NickFect family were designed to condense nucleic acid to nanoparticles and thereby protect it against degradation by nucleases, as well as to promote cellular uptake and endosomal escape of formed nanoparticles. Here we show that in bloodstream, CPP-NA nanoparticles acquire a novel nano-biomolecular interface and recruit into protein corona anti-opsonising proteins like serum albumin, clusterin and apolipoproteins at higher extent compared to only nucleic acids-containing particles. Moreover, CPPs reduce inclusion of C3 protein in protein corona on pDNA particles and thereby might prolong circulation of respective nanoparticles in organism before elimination (Fig. 3c, d).^83,84^

### Protein corona on CPP-SCO nanoparticles controls the efficiency of internalisation and splicing switching by SCO in cells

The divergence of protein corona composition in sera of different animals prompted us to assess the uptake of PF14-SCO nanoparticles carrying dissimilar protein corona by cultured cells. The association of PF14 nanoparticles formed with fluorescently labelled (Cy5) SCO with HeLa cells varied markedly in the culture media containing sera of different animals. The protein corona acquired in presence of bovine serum induced the highest internalization of PF14-SCO-Cy5 complexes (Fig. 5a, e), whereas the presence of murine or human serum led to lower association of nanoparticles with HeLa cells (Fig. 5b, c). In concordance with TEM experiments, fluorescence microscopy also revealed formation of larger clusters of PF14-SCO-Cy5 complexes on the cell surface in the presence of sera, compared to protein-free culture medium, suggesting association of complexes into agglomerates in these conditions. PF14-SCO-Cy5 complexes were not targeted to acidic organelles during the first hour of internalisation, suggesting that mostly macropinocytosis and caveolae-mediated endocytosis, i.e. the cellular uptake pathways typically triggered by PF14, were involved.^89^ Surprisingly, the intensity of Lysosensor signal varied substantially in cells grown in media containing different sera. However, the lowest Lysosensor signal in human serum-containing growth media, and low internalisation efficiency of nanoparticles cannot be caused by impaired viability of cells, since HeLa cells are known to proliferate in the presence of human serum in normal manner and we did not detect any changes in viability.^90^

As mentioned earlier, from each serum a specific subset of proteins was recruited into protein corona. Interestingly, the most efficient internalisation of PF14-SCO-Cy5 nanoparticles was observed in bovine sera containing culture medium, where forming PC contained fetuins that are known to promote the cellular uptake of these nanocomplexes.^52,91^

**Figure 5.**
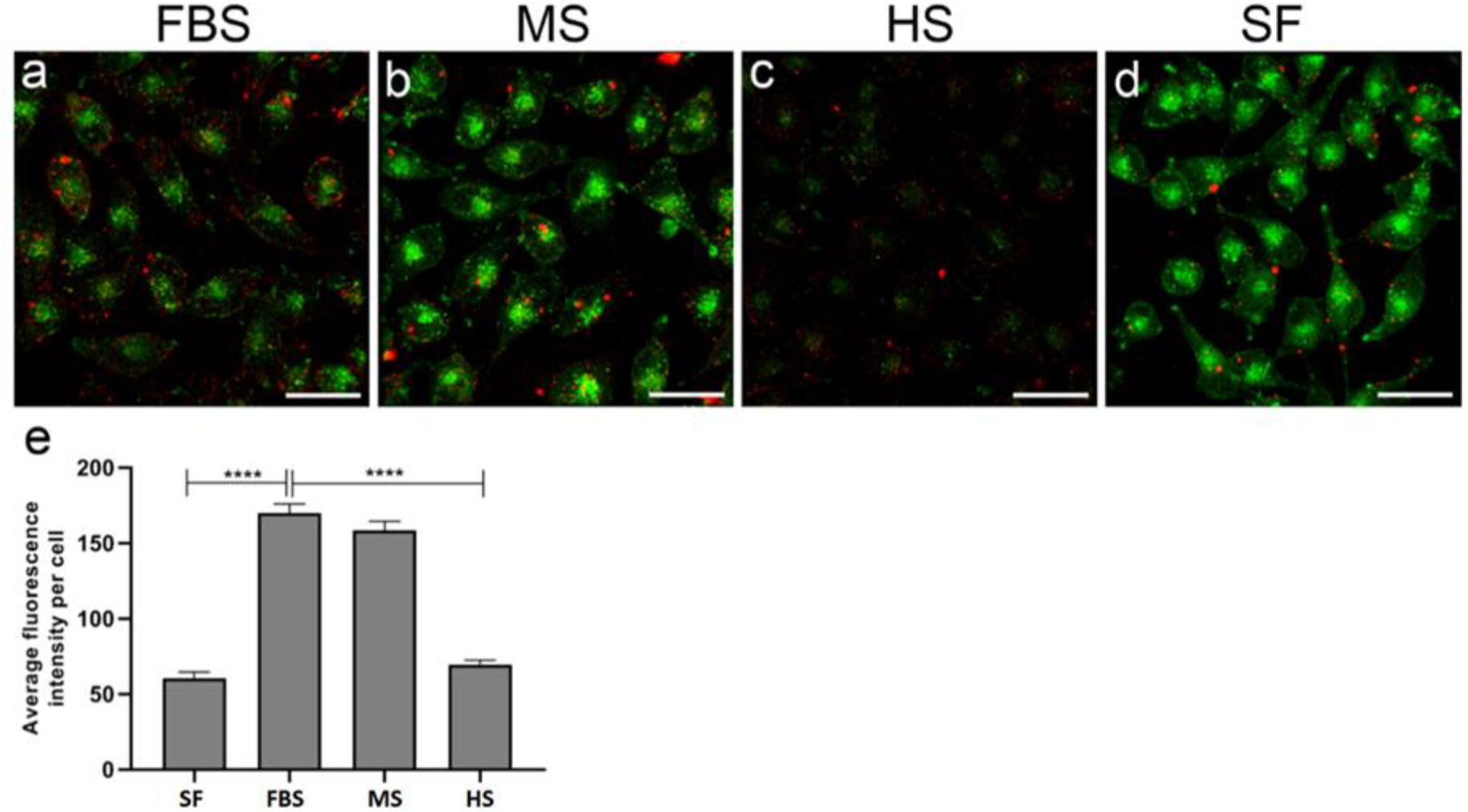
Association of PF14-SCO nanoparticles carrying different protein corona with cells. Complexes of PF14 with Cy5-SCO prepared at molar ratio 5, were incubated with HeLa cells in culture medium containing 10% of foetal bovine serum (**a, FBS**), murine serum (**b, MS**), or human serum (**c, HS**), or lacking serum (**d, SF**) for 1 h. Lysosomes and other acidic organelles were visualised with Lysosensor Green. Localisation of nanoparticles based on Cy5 label on SCO (red), and acidic organelles (green) was captured in living cells with confocal microscope FV1000 and cell-associated SCO-Cy5 was quantified using Autoquant software (**e**). Scale bar 30 μm. Representative images from three independent experiments are presented. In panel **e**, bars represent average fluorescence intensity per cell ±SEM. Data was analysed using one-way ANOVA with post-hoc Tukey’s test. ****p < 0.0001.

Unequal uptake of PF14/SCO nanoparticles having different protein corona by HeLa cells also suggests that the effect of SCO cargo inside cells may also differ. In concordance with the differences in internalisation efficiency of nanoparticles, also correction of mRNA splicing by delivered SCO in reporter cell-line HeLa pLuc705 varied substantially in culture media containing different sera. The increase in expression/activity of luciferase was the highest in calf serum containing media exceeding about 2-fold the activity in cells treated with human-protein corona carrying nanoparticles (Fig. 6). These results are also in good correlation with different cellular uptake of SCO in different sera (Fig. 5)

**Figure 6.**
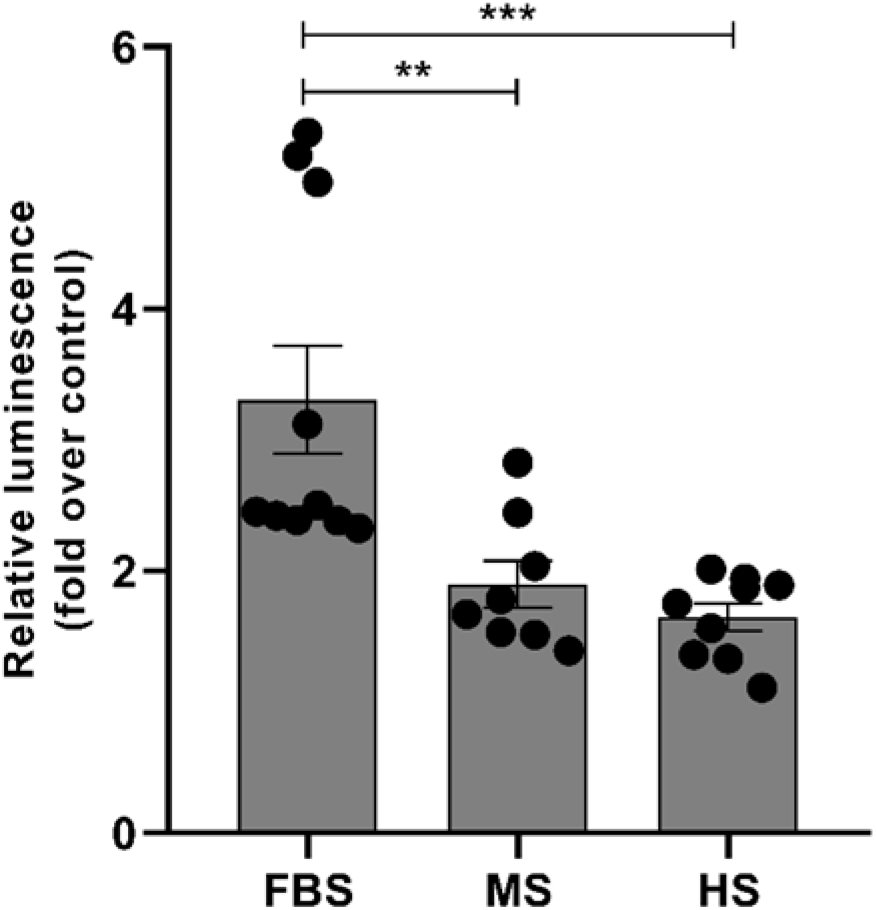
Splicing switching induced by PF14-SCO nanoparticles carrying different protein corona. HeLa pLuc705 cells were incubated with nanoparticles of 500 nM PF14 with 100 nM SCO in culture media containing 10% of foetal bovine serum (FBS), murine serum (MS) or human serum (HS) for 4 h and luminescence was quantified after additional 20 h of incubation. Data was analysed using one-way ANOVA with post-hoc Tukey’s test. Each dataset is presented as mean ± SEM. *p < 0.05, ***p < 0.0005.

## Conclusions

In the development of CPP-nucleic acid nanoparticles for biomedical use, it is important to keep in mind that after their contact with biological fluids, the particles are rapidly coated with a protein corona that might significantly alter the fate and efficacy of the nanoparticles. Our study demonstrates that major components of protein corona that forms on the CPP-nucleic acid nanoparticles in animal sera and plasma are proteins involved in transport of various substances, haemostasis, and complement system. Most importantly, formation of protein corona, as well as its content is species-specific and is dependent on the properties of both, nucleic acid and condensing CPP. We demonstrated that variations in the protein corona composition in different sera lead to unequal association of nanoparticles with cells and dissimilar biological response to delivered nucleic acid molecules inside cells. This might be one of the main reasons, why CPP-nucleic acid nanoparticles that are highly efficient *in vitro*,often fail to yield expected outcome *in vivo* upon systemic administration. Thus, for CPP-mediated transport, transition from bench to bedside, still remains complicated, unless topical delivery is used where less complex protein corona forms around nanoparticles.^11,96,97^

## Materials and Methods

PF14 (see sequences and characteristics of used CPPs and nucleic acids in Table 1), C22-PF14 and NF55 were purchased from Pepmic (China). Splicing correcting oligonucleotides with 2’OMe modification and phosphorothioate backbone, and labelled with Cy5 or biotin were from Microsynth (Switzerland), or Metabion (Germany), and pGL3 plasmid was from Promega, (4.7 kbp, USA). All other reagents were from Sigma-Aldrich (Merck KGaA, Germany), if not stated otherwise.

### Plasma and serum collection

For preparation of murine plasma, 8 weeks old mice (strain C57Bl/6, University of Tartu, IMCB Animal research facility) were used. 3.2% (w/v) sodium citrate in a total volume of body weight (gr) was administered intravenously in the vena cava 20–30 seconds prior to blood drawing from the same vein into a syringe. Blood samples were centrifuged for 15 min at 3000 rpm at room temperature. For removal of remaining cells and platelets, plasma was centrifuged for 5 min at 2000 × g, aliquoted, and immediately frozen at −25 °C.

For preparation of murine serum, mice were decapitated; blood was collected and allowed to clot at room temperature for 20 min. After centrifugation (10 min at 2000 × g), upper fraction was removed, aliquoted, and immediately frozen at −25 °C.

Foetal bovine serum (FBS) was purchased from Corning, Life Sciences (USA), human serum from Sigma-Aldrich, and standard human plasma from Dade Behring (USA).

### Labelling of pDNA with biotin

Biotin was covalently coupled to pDNA (pGL3 plasmid) by using Nucleic Acid Labeling Kit according to manufacturers’ protocol (*Label* IT^®^ Nucleic Acid Labeling Kit, Biotin, Mirus Bio, USA). Briefly, the biotin reagent dissolved in Reconstitution Solution was mixed with pDNA (1 μg/μl) at a ratio 1:5 (v/v) in Labeling Buffer A for 1 h at 37 °C. The labelled plasmid was separated from free biotin by ethanol precipitation, dissolved in Milli-Q (MQ) water and stored as aliquots at −25 °C.^49^

### Nanoparticle formation and pull-down

Pierce™ Streptavidin superparamagnetic beads (Thermo Fisher Scientific) were washed with 0.1% Tween-20 in TBS before incubation with 1 μg of biotinylated pGL3 or 200 nM biotinylated SCO in MQ water in 1/10th of the final volume for 15 min at 37 °C with mild shaking (300 rpm). For nanoparticle formation, CPPs were added at charge ratio (CR, N/P ratio) 2 (CPP:pGL3) or molar ratio (MR) 5 (CPP:SCO) and incubated for 30 min at room temperature with mild shaking. After addition of respective serum or plasma, streptavidin-beads were incubated for 5 or 60 min at room temperature with mild shaking. Then, streptavidin-beads were washed 3 times with PBS (5 min each) or TBS (for mass spectrometry analysis) and beads were collected on a magnetic stand.

### Dynamic light scattering (DLS) and zeta potential measurements

Hydrodynamic mean diameter and zeta potential of the nanoparticles were measured using a Zetasizer Nano ZS apparatus (Malvern Instruments, UK). CPP-nucleic acid nanoparticles were formed in MQ water at MR 5, for SCO-CPP complexes, and at CR 2, for pDNA-CPP complexes at room temperature for 30 min. For the hydrodynamic diameter and zeta-potential measurement, nanoparticles solutions were diluted to reach 0.5 μM SCO, 0.2 μg/ml pGL3 concentration in MQ water or 1 mM NaCl, respectively. Mean hydrodynamic diameter values are represented as Z-average (± SD), based on four measurements for each nanoparticles, and zeta potential values represent arithmetic mean of four measurements.

### Protein separation and silver staining

2× SDS loading buffer was added to complexes on beads and samples were denaturated at 96 °C for 5 min. Proteins were separated by electrophoresis on a 10% SDS polyacrylamide gel and visualised with Silver Stain Plus Kit (Bio-Rad Laboratories AB, Sweden) according to manufacturer’s modified silver stain protocol.

### Western blot

2× SDS sample buffer was added to samples, sonicated three times at maximum intensity (Bioruptor^®^, Diagenode, Belgium) and the samples were denaturated at 65 °C for 15 min. Proteins were separated by electrophoresis on a 9% SDS polyacrylamide gel and transferred to polyvinylidene difluoride (PVDF) membrane. Sites of nonspecific binding were saturated with 5% non-fat dry milk (NFDM) in TBS for 1 h. After washing with TBS, the membranes were incubated with rabbit monoclonal anti-C3 antibody (1:2000; EPR19394, Abcam, UK) in 0.5% NFDM in TBS at 4 °C overnight, washed three times (5 min each) and incubated with alkaline phosphatase anti-rabbit IgG (1:2000 dilution in 0.5% NFDM in TBS; Abcam, UK) at room temperature for 1 h. The signal was developed with Nitro Blue tetrazolium and 5-bromo-4-chloroindol-2-yl phosphate in the alkaline phosphatase buffer (100 mM NaCl, 10 mM MgCl_2_, 100 mM Tris–HCl, pH 9.5). Quantification of Western blot images (SFig. 1-4) was carried out using ChemiDoc™ XRS+ Image Lab™ Software (Bio-Rad Laboratories AB, Sweden); statistical analysis was performed with GraphPad Prism software (version 5.0, GraphPad, USA).

### Mass-spectrometry

CPP-pDNA nanoparticles or b-pGL3 with associated proteins were captured on streptavidin-coated superparamagnetic beads, washed and collected as above, and subjected to trypsin digestion. Subsequent mass-spectrometry analysis was performed at the Core facility for Proteomics of University of Tartu as described earlier.^92^ Briefly, the peptide mixtures were separated on 3 μm ReproSil-Pur C18AQ 15 cm × 75 μm ID emitter-column (New Objective) using Agilent 1200 series nano-LC, and detected with an LTQ Orbitrap XL (Thermo Fisher Scientific) mass-spectrometer. Mass-spectrometric raw data were analysed with MaxQuant 1.5.3.172. Overall number of identified peptides of a protein group is presented. Data represents three independent experiments with three technical replicates.

### Electron microscopy of nanoparticles

For analysis of nanoparticles’ morphology and protein corona, splice-correcting oligonucleotide (SCO) was condensed to nanoparticles with PF14, and labelled with colloidal gold tag. Biotin-tagged SCO (b-SCO) at 2 μM concentration was complexed with 10 μM PF14 (MR 5) in MQ water at room temperature for 30 min.^50^ For negative staining, 10 μl of sample was absorbed to copper grids covered with Formvar film and carbon layer for 2 min, and excess of solution was gently removed. Next, grids were exposed to 2% aqueous uranyl acetate for 1 min. After removing excess of stain with filter paper, specimens were dried in air.

For visualisation of protein corona, b-SCO was first coupled to neutravidin (labelled with 10 nm colloidal gold)^93^ and then complexed with PF14. The formed NPs were exposed to 50% bovine, murine or human serum^94^ in Dulbecco’s modified Eagle’s medium (DMEM, Life Sciences, Thermo Fisher Scientific, Germany) for 30 min, and absorbed to TEM grids. Afterwards grids were washed three times with MQ water and air-dried. The specimens were examined at 120 kV accelerating voltage with FEI Tecnai G2 Spirit transmission electron microscope (FEI, The Netherlands).

### Translocation of PF14-SCO nanoparticles into cells in different sera containing media and splicing correction by SCO

HeLa pLuc705 cells,^95^ kindly provided by Prof. R. Kole, were grown in DMEM supplemented with 10% FBS and antibiotics in humidified atmosphere containing 5% CO_2_ at 37 °C. For visualisation of acidic organelles, the medium on cells was exchanged to serum-free culture medium that contained 1 μM Lysosensor Green DND-189 (Molecular probes Inc.). After incubation for 1 h at 37 °C, it was replaced with PF14-SCO-Cy5 nanoparticles containing media. For that, PF14 and SCO-Cy5 complexes were formed in MQ water at MR 5 as described above, incubated at room temperature for 30 min^89^ and diluted with DMEM containing 10% bovine, murine or human serum. PF14 and SCO-Cy5 nanoparticles were applied to cells to reach 1 μM and 200 nM concentrations, respectively. After incubation for 1 h with complexes, cells were washed with PBS and analysed with confocal laser scanning microscope (FV1000, Olympus, USA). The amount of cell-associated nanoparticles was quantified by Cy5 signal using Autoquant X3 software.

For measurement of luciferase activity, 7 000 cells per well were seeded onto 96-well plate 24 h before transfection. Next day, PF14-SCO complexes of 500 nM PF14 and 100 nM SCO (applied as triplicates) were added to the cells in DMEM that contained 10% bovine, murine or human serum. After incubation for 4 h, the complex-containing solutions were replaced by medium with 10% FBS and luciferase activity was measured 20 h later. Before the measurement, cells were washed twice with PBS, and lysed with Cell Culture Lysis Reagent (Promega, USA) using one freeze-thaw cycle. Luminescence was quantified in relative light units (RLU) using Promega’s luciferase assay system and microplate luminometer (Infinite 200 M, Tecan, Austria).^52^ The luciferase activity data was normalised to protein content measured by DC Protein determination kit (Bio-Rad Laboratories, Inc., USA) and SCO effect was expressed as a fold-change over untreated cells. The effect of protein corona was expressed as the ratio of luciferase activity of the cells treated with PF14-SCO nanoparticles and control cells.

### Statistical analyses

All statistical analyses were performed using Prism 5.0 software (GraphPad Software). Results of at least three independent experiments are presented ±SEM. Statistical significance of differences was analysed by one-way ANOVA with post-hoc Tukey’s test at a significance level of 0.05. Asterisks indicate a statistically significant difference between two groups.

## Abbreviations

C3: component 3 of complement system
C22-PF14: PepFect 14 analogue with a 22 carbon-fatty acid (behenyl)
CPP: cell-penetrating peptide
MS: mass-spectrometry
NF55: NickFect 55 peptide
pDNA: plasmid DNA
PF14: PepFect 14 peptide
SCO: splice-correcting oligonucleotide.

## Acknowledgments

We thank Kaida Koppel for preparation of TEM specimens, Rahel Paloots and Carmen Tali for initial characterisation of PC components. We highly appreciate help of Sulev Kuuse for preparation of murine blood serum and plasma. This work was supported by the Estonian Ministry of Education and Research (0180019s11), Estonian Research Council (PUT1617P, PRG1169), Institute of Technology basic financing grant (PLTTI20912) to MP, and Leo Foundation (LF17040).

## Author contributions

**Annely Lorents:** conceptualization, investigation, formal analysis, methodology, validation, visualisation, writing – original draft. **Maria Maloverjan**: investigation, validation, visualisation, writing. **Kärt Padari**: investigation, methodology, validation, visualisation, writing. **Margus Pooga**: conceptualization, methodology, funding acquisition, project administration, supervision, validation, writing – original draft, review & editing.

